# Covert spatial attention speeds target individuation

**DOI:** 10.1101/838557

**Authors:** Joshua J. Foster, Emma M. Bsales, Edward Awh

## Abstract

Covert spatial attention has long been thought to speed visual processing. Psychophysics studies have shown that target information accrues faster at attended locations than at unattended locations. However, with behavioral evidence alone, it is difficult to determine whether attention speeds visual processing of the target, or subsequent post-perceptual stages of processing (e.g. decision making and response selection). Moreover, while many studies have shown that that attention can boost the *amplitude* of visually-evoked neural responses, no effect has been observed on the *latency* of those neural responses. Here, we offer new evidence that may reconcile the neural and behavioral findings. Our study focused on the N2pc, an EEG marker of visual selection that has been linked with object individuation – the formation of an object representation that is distinct from the background and from other objects. In two experiments, we manipulated whether or not covert attention was precisely deployed to the location of an impending search target. We found that the target-evoked N2pc onset approximately 20 ms earlier when the target location was cued than when it was not cued. Thus, although attention may not speed the earliest stages of sensory processing, attention does speed the critical transition between raw sensory encoding and the formation of individuated object representations.

**Significance Statement:** Covert spatial attention improves processing at attended locations. Past behavioral studies have shown that information about visual targets accrues faster at attended than at unattended locations. However, it has remained unclear whether attention speeds perceptual analysis or subsequent post-perceptual stages of processing. Here we present robust evidence that attention speeds the N2pc, an electrophysiological signal that indexes the formation of individuated object representations. Our findings show that attention speeds a relatively early stage of perceptual processing, while also elucidating the specific perceptual process that is speeded.

## Introduction

Covert spatial attention is thought to improve the fidelity and speed of visual processing (1–4). Although there is strong evidence that attention improves the fidelity of processing (1, 2), evidence that attention speeds visual processing has been harder to come by. The strongest evidence that attention speeds visual processing comes from psychophysics studies that have obtained independent measures of processing speed and visual discriminability to show that target information accrues *faster* at attended relative to unattended locations (5–8). However, because this work relies on behavioral responses, it is difficult to establish conclusively whether attention speeds perceptual analysis of the target, or subsequent post-perceptual stages of processing (e.g., decision making or response selection). Adding to this uncertainty, a broad body of EEG work has revealed that attention influences the *amplitude* but not the *latency* of early visually evoked neural responses (9–12). Likewise, studies in non-human primates have reported either no effect of covert attention on the latency of responses in visual cortex (13, 14) or have found very small differences of 1-2 ms (15), which are an order of magnitude smaller than the effects on processing speed estimated from behavior [e.g. Ref (5)]. Thus, neural and behavioral data have not yet converged on a common answer to the fundamental question of whether attention speeds early visual processing.

In the present work, we examined how covert spatial attention affects the N2pc, a visually-evoked EEG signal that has long been viewed as a neural marker of visual selection (16, 17). The N2pc is a transient negative deflection that is seen contralateral to a selected item in a visual display, typically occurring 150-300 ms after display onset. Prior work has shown that N2pc latency is sensitive to a range of manipulations such as search difficulty (18), temporal cues (19), and reward (20). This work suggests that the N2pc may be particularly well suited for testing whether spatial attention alters the speed of visual processes. First, however, it is important to consider what aspect of visual processing is indexed by the N2pc.

Although the N2pc was initially thought to index a shift of covert attention (16, 17, 21), recent studies suggest that it may instead track a process that occurs *after* the initial shift of attention. For example, Kiss and colleagues (22) showed that the amplitude of the N2pc evoked by a target during visual search was equivalent when a cue indicated the target’s hemifield in advance – enabling a preparatory shift of attention – compared to when an uninformative cue was presented. Thus, Kiss et al. concluded that the N2pc does not index shifts of attention *per se*. An emerging view is that the N2pc indexes *object individuation* (23, 24), the formation of an object representation that is segregated from the background and from other items in the display (25, 26). In support of this view, N2pc amplitude increases monotonically with the number of items that are individuated during rapid enumeration tasks (27, 28), and tracks individual differences in tasks that critically depend on object individuation, such as target enumeration (24) and multiple object tracking (29). Moreover, the N2pc does not track the number of targets in a display when the task does not require that the targets are individuated (e.g. target detection rather than enumeration) (27). Thus, the N2pc appears to index a volitionally controlled process of object individuation rather than shifts of spatial attention.

In this context, the current work provides two theoretical advances. First, we verified the conclusion of Kiss and colleagues (22) while addressing an important caveat to their conclusions. Specifically, observers in the Kiss et al. study were cued to the hemifield rather than the precise position of the target, leaving open the possibility that N2pc amplitude in the cued condition reflected the re-focusing of attention within the cued hemifield. By contrast, in the current study we cued the precise position of the target, and we used the topography of alpha-band (8–12 Hz) oscillations to track the locus of covert attention (30, 31), thereby verifying that observers attended the cued location prior to the target onset. Having verified that attention was precisely deployed to the target position following informative cues, we observed equivalent N2pc amplitudes following informative and uninformative cues, confirming Kiss and colleagues (22) conclusion that the N2pc component does not track shifts of attention *per se*. Instead, our results support the view that the N2pc reflects individuation of a visual target because the target must still be individuated when spatial attention is already focused at the target location. However, the key theoretical advance in the present work is that attention speeds the perceptual process of object individuation. In two studies we found that the N2pc component occurred approximately 20 ms earlier when the location of the search target was cued in advance, suggesting that the search target was individuated earlier when attention was already focused at the target location. Thus, our findings point to a reconciliation of past behavioral and neural studies of how attention influences the latency of visual processing. While spatial attention may not change the latency of very early sensory responses, our findings suggest that attention does speed the transformation from raw sensory input to discrete representations of individuated objects.

## Results

In two experiments, we tested whether spatial cues in advance of a search array influence the latency of the N2pc component. Observers performed a cued-search task (Fig. 1). On each trial, observers searched for a target – a diamond among squares – and reported whether the target was missing its left or right corner [also see (22)]. We used a spatial cue to manipulate whether or not covert attention was focused at the target location prior to the onset of the search array. In some blocks, the cue indicated the precise location of the impending target (*informative* cue), which allowed observers to focus covert attention on the target location in advance. In other blocks, the cue was spatially uninformative (*uninformative* cue), such that observers needed to monitor all eight positions for the target.

**Figure 1.**
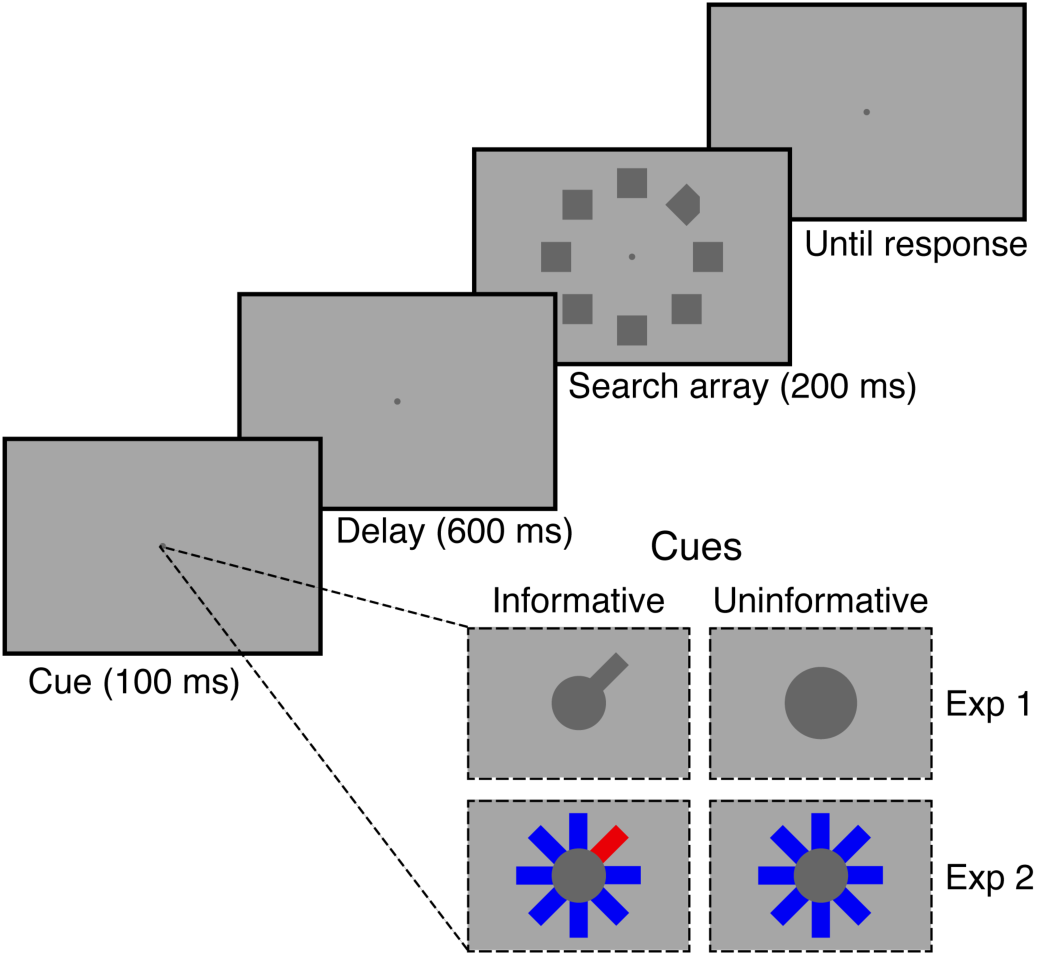
Cued-search task. Observers searched for a diamond among squares, and reported which corner was missing (left or right). The target could appear in any of eight positions around the fixation point. The search array was preceded by a cue. In some blocks, the cue indicated the exact position of the upcoming target (informative cue). In other blocks, the cue was spatially uninformative (uninformative cue).

Cueing the position of the target improved search performance in both experiments. Figure 2 shows median response times (RTs) and accuracy (% correct) as a function of cue type (informative vs. uninformative). In Experiment 1, median RTs were 45 ms faster on average following informative cues (*M* = 503 ms, *SD* = 46) than following neutral cues (*M* = 548 ms, *SD* = 39), *t*(15) = 7.49, *p* < .001, and accuracy was higher following informative cues (*M* = 97.7%, *SD* = 1.3) than following neutral cues (*M* = 95.5%, *SD* = 2.7), *t*(15) = −4.94, *p* < .001. We replicated this pattern in Experiment 2: median RTs were 54 ms faster on average following informative cues (*M* = 496 ms, *SD* = 62) than following neutral cues (*M* 550 ms, *SD* = 73), *t*(25) = 12.79, *p* < .001, and accuracy was higher following informative cues (*M* = 96.9%, *SD* = 2.3) than following neutral cues (*M* = 95.2%, *SD* = 2.9), *t*(25) = −5.58, *p* < .001. These results show that observers made use of informative cues, attending the target location in advance. Below, this conclusion will be corroborated by our analysis of alpha activity during the time window between cue and target onset.

**Figure 2.**
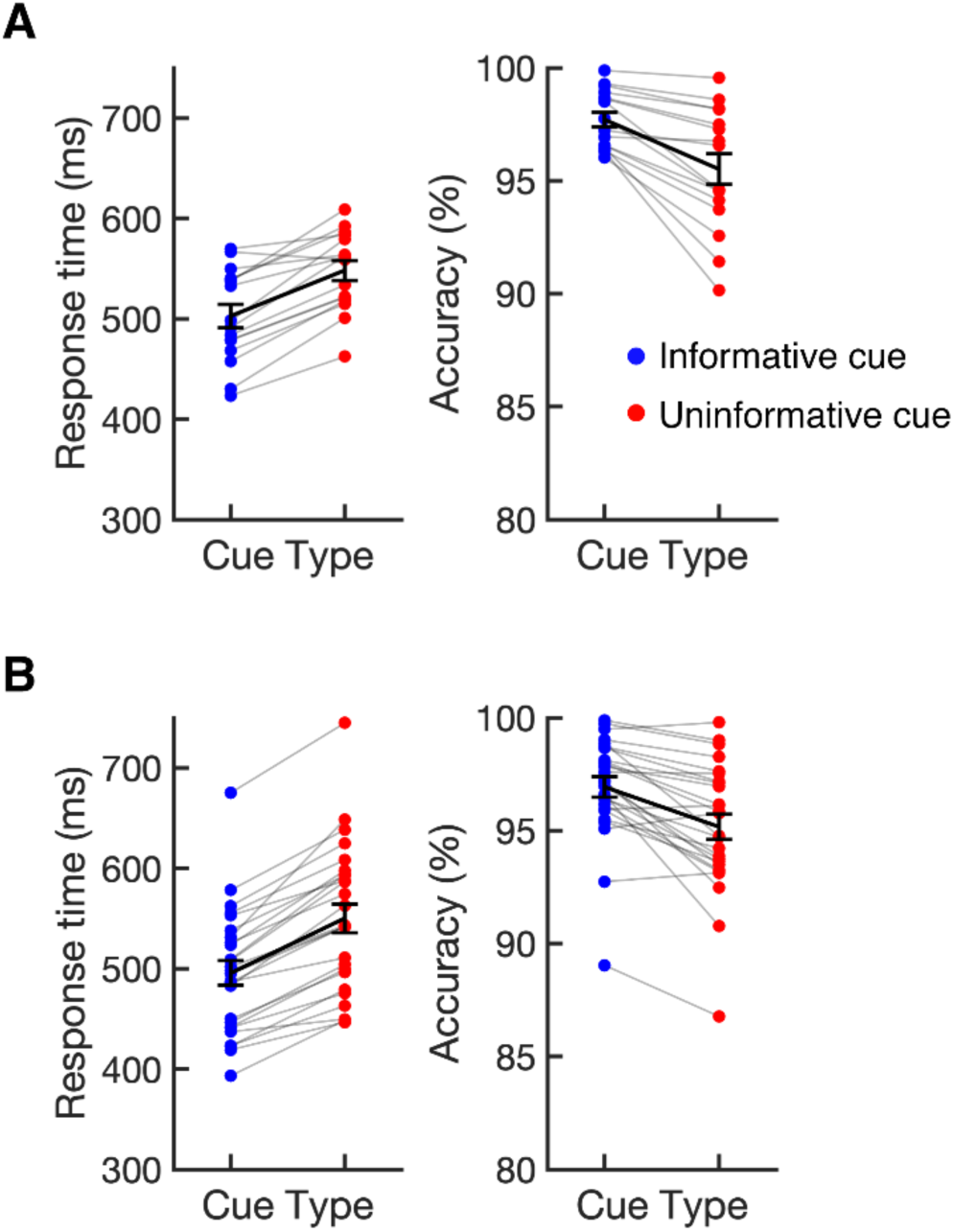
Median response times (RTs) and accuracy (% correct) following informative and uninformative cues for Experiment 1 (A) and Experiment 2 (B). Dots linked by grey lines show individual participants. The black lines show the sample mean. Error bars show ±1 SEM across subjects.

### Covert spatial attention speeded the N2pc component

To test whether covert attention speeds visual processing, we tested whether our manipulation of covert attention (informative vs. uninformative cues) influenced the latency of the target-evoked N2pc seen following the onset of the search array. Figure 3 the shows the contralateral – ipsilateral difference waves locked to the onset of the search array for both experiments. The target-evoked N2pc is the negative deflection in the difference wave occurring between 150 and 300 ms after onset of the search array. We observed a robust N2pc following both informative and neutral cues. We measured the onset of the N2pc as the time at which the N2pc reached 50% of its maximum amplitude, and used a jackknife procedure to test whether N2pc latency differed between informative and uninformative cue conditions (see Materials and Methods). In Experiment 1, we found that the target-evoked N2pc onset 18 ms earlier following informative cues than following uninformative cues (Fig. 3a), *t*(15) = 2.19, *p* < .05 (one-tailed test). In Experiment 2, we sought to replicate this effect with a larger sample size to increase statistical power (see Materials and Methods). In Experiment 2, we again found that the N2pc earlier (this time 22 ms earlier) following informative cues than following uninformative cues (Fig 3b), *t*(25) = 3.82, *p* < .001 (one-tailed test). Control analyses confirmed that the effect of cueing on the latency of the N2pc cannot be explained by very small residual biases in gaze toward the cued location that remained after our stringent artifact rejection procedures (see Supplemental Information). Together, these results provide clear evidence that covert attention speeds the N2pc.

**Figure 3.**
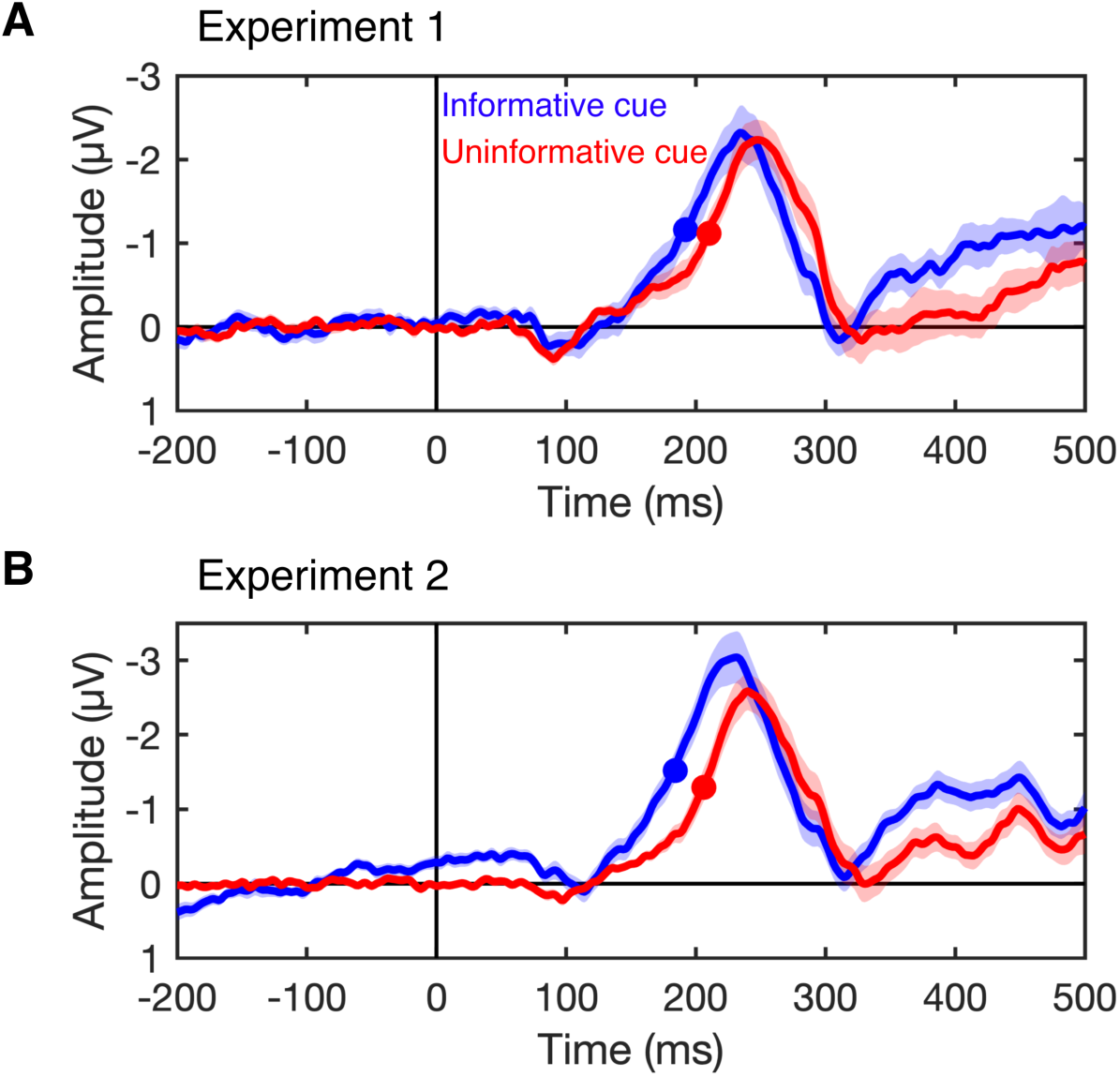
N2pc difference waves for Experiment 1 (A) and Experiment 2 (B). The plots show the N2pc difference waves (contralateral minus ipsilateral) time locked to the onset of the search array following informative cues (blue) and uninformative cues (red). The N2pc is the negative deflection seen between 150 and 300 ms (note that negative is plotted up). The filled circles mark the time at which the N2pc reached the onset criterion (50% of maximum amplitude). The N2pc onset earlier following informative cues than following uninformative cues. Shaded error bars show ±1 SEM across subjects.

### The N2pc does not index shifts of attention

The N2pc is often taken as an index of a shift of spatial attention to a target stimulus (16, 21). However, Kiss and colleagues (22), using a cued-search paradigm similar to ours, found that the amplitude of the N2pc evoked by a target during visual search was equivalent following informative and uninformative cues. Based on this finding, they argued that the N2pc does to index a shift of spatial attention because observers had shifted attention to the cued location prior to the search array following informative cues. However, Kiss and colleagues cued the hemifield (left or right) that the target would appear in rather than its exact position. Thus, the cued enabled observers to attend the cued side of space, but not the exact position of the target, prior to the search array. As a result, observers likely broadly attended the cued side of space following the cue, before focusing attention on the exact location of the target once the search array appeared. The N2pc that Kiss and colleagues observed following informative cues may have reflected this refocusing of spatial attention within the target’s hemifield. By contrast, informative cues in our experiments indicated exactly where the target would appear, allowing observers to precisely attend the target location in advance. To verify that observers in our experiments precisely attended the cued location prior to the search array, we examined alpha-band (8-12 Hz) oscillations that precisely track when and where spatial attention is deployed (31–33).

To this end, we used an inverted encoding model [IEM (31, 34–36)] to reconstruct channel-tuning functions (CTFs) from the pattern of alpha across the scalp that track the allocation of spatial attention (31). This approach assumes that alpha power at each electrode reflects the joint activity of a number of spatially tuned channels (or neuronal populations). By first estimating the contributions of these channels to activity measured at each electrode on the scalp, the model can then be inverted so that the underlying response of these spatial channels can be estimated from the pattern of alpha power across the scalp (31, 37). We used data from the informative-cue condition to train the model (i.e., estimate the contribution of each spatial channel to each electrode), and then inverted the model to reconstruct the profile of activity across the spatially selective channels for each of the conditions separately (see Materials and Methods). The resulting alpha CTFs reflect the spatial selectivity of population-level alpha activity measured with EEG (31, 37).

Alpha activity precisely tracked the cued location. Figure 4 shows the channel responses as a function of the impending target location following informative cues (left) and uninformative cues (right). Because filtering alpha-band activity leads to temporal smearing, we focused on a window 300-500 ms after cue onset (ending 200 ms prior to the onset of the search array). Channel responses in this window purely reflect activity prior to the onset of the search array (see Materials and Methods). Following informative cues, we found that the peak channel response tracked the cued target location, with the peak channel response always in the channel tuned for the cued target location, or in the neighboring channel. As expected, the peak channel response did not track the target location (which was not cued) following uninformative cues. Figure 5 shows the time course of spatially selective of alpha-band CTF (measured as CTF slope, see Materials and Methods). Higher CTF selectivity indicate stronger tuning for the cued/target location. Following informative cues, we found that spatially tuned alpha activity emerged shortly after cue onset, and persisted throughout the search array. Following uninformative cues, spatially tuned alpha activity did not emerge until after the onset of the search array.

**Figure 4.**
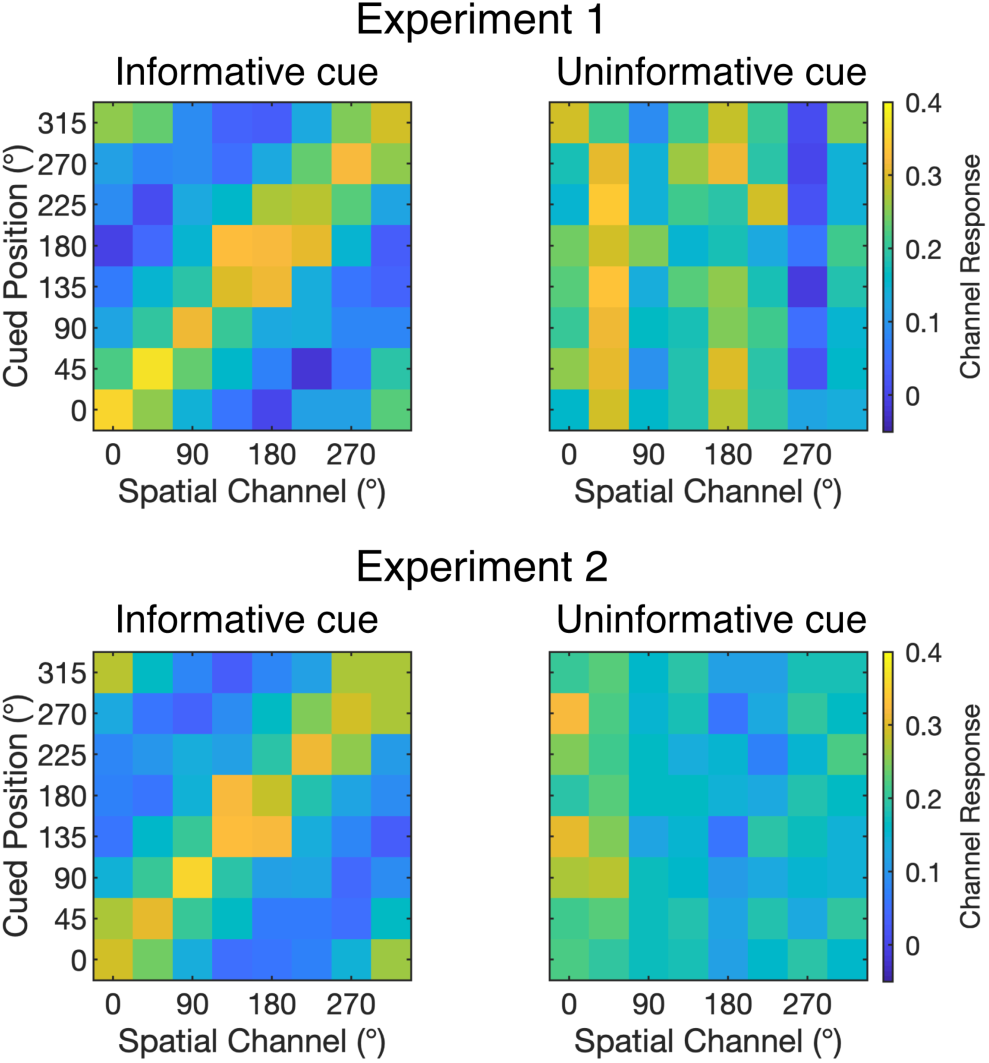
Reconstructed responses of spatially tuned channels as a function of cued position 300-500 ms after cue onset following informative and uninformative cues in each experiment. We found that the peak channel response tracked the location of the impending target following informative cues (left) but not following uninformative cues (right).

**Figure 5.**
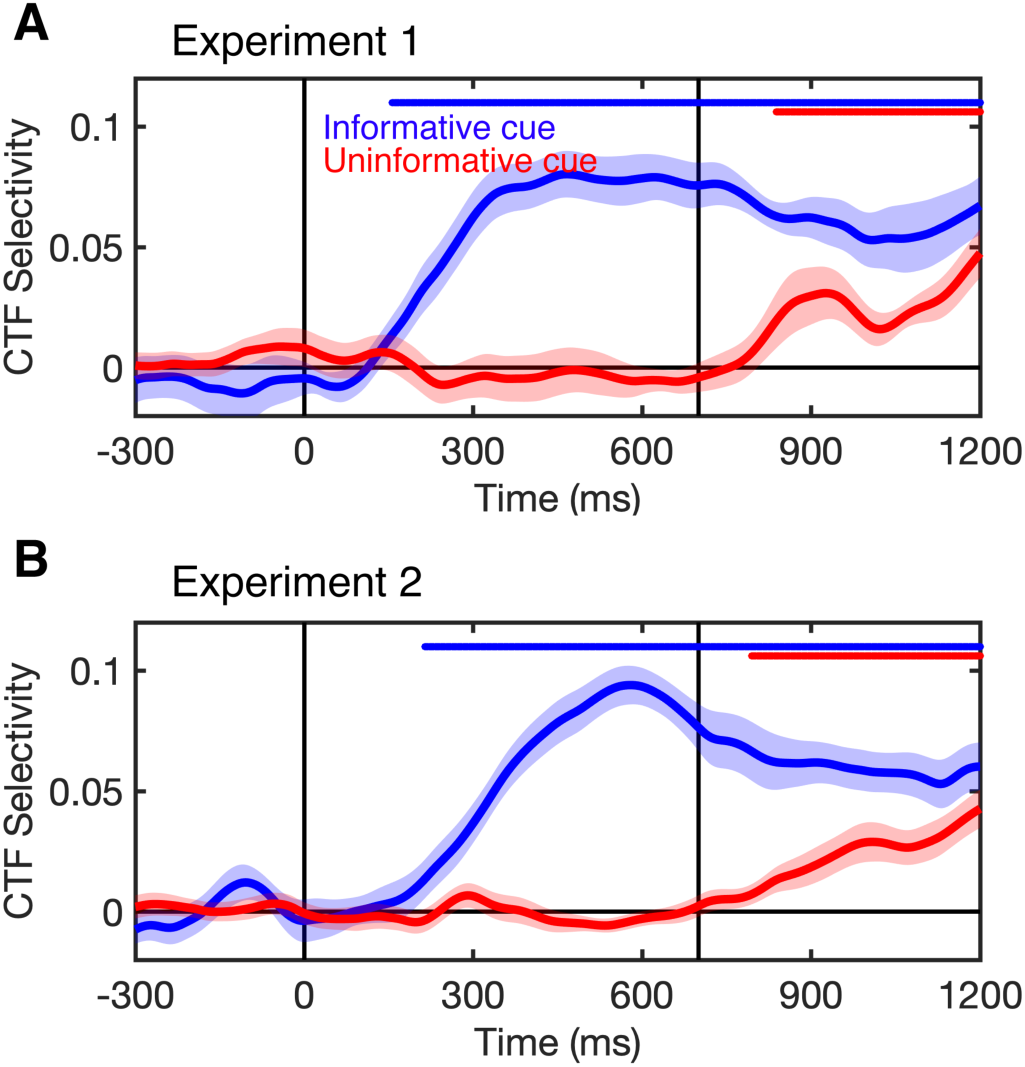
The spatial selectivity of alpha-band channel-tuning functions (CTFs) across time (measured as CTF slope, see Materials and Methods) following informative (blue) and uninformative (red) cues. In both experiments, we found that spatially selective activity that tracked the cued location emerged shortly after cue onset (0 ms) following informative cues, but not until after the search array appeared (700 ms) following uninformative cues. The blue (informative cue) and red (uninformative cue) markers at the top of each panel indicate periods of above-chance selectivity obtained using a cluster-based permutation test (Materials and Methods). Shaded error bars show ±1 SEM across subjects.

Taken together, these findings provide evidence that observers precisely attended the cued locations prior to the search display following informative cues. Thus, we found a robust N2pc following informative cues (Fig. 3) even when observers had attended the target location prior to the search array. These findings provide strong support for the Kiss et al. conclusion that the N2pc does not index shifts of attention *per se*. Instead, our results support the view that the N2pc indexes object individuation (23, 24) – the formation of an object representation that is segregated from the background and from other items in the display (25, 26). The target stimulus can only be individuated once a search array has been presented. Therefore, the individuation account of the N2pc predicts that a target-evoked N2pc should be seen following target onset, even when covert attention is already focused at the target location.

## Discussion

The idea that covert spatial attention speeds visual processing has a long history (3, 4, 38). In recent years, psychophysics has supported this idea, suggesting that information about a visual target accrues faster from attended locations than from unattended locations (5–8). However, because this work relies on behavioral responses, it is difficult to tell whether attention speeds the perceptual processing of the target, or enables more efficient use of this information in subsequent post-perceptual stages of processing (e.g., decision making and response selection). At the same time, work focused on the latency of target-evoked neural responses has challenged the claim that sensory processing is speeded by attention, showing increased amplitude of early visual responses to attended stimuli, but little or no effect on the latency of these responses (9–15). Here, our findings offer a reconciliation of these different lines of work.

We provide clear evidence that attention does in fact speed visually evoked responses. Observers performed a cued-search task. In some blocks, a spatial cue indicated the exact position of the upcoming target, allowing observers to focus spatial attention at the relevant location. In other blocks, the cue was uninformative. We examined how this manipulation of spatial attention influenced the N2pc, a negative deflection seen at posterior sites contralateral to visually selected stimuli. In two experiments, we found that the N2pc occurred earlier following informative relative to uninformative cues. In light of evidence that the N2pc indexes the formation of individuated object representations (23, 24, 27, 28), our findings suggest that spatial attention speeds this aspect of object perception.

Thus, our findings reconcile the apparent conflict between evidence from psychophysics that attention speeds perceptual processing and neural measurements that have found that attention does not substantially alter the latency of visually evoked responses. We propose that attention does not speed the first feedforward sweep of visually evoked activity, but does speed the subsequent formation of individuated object representations. Although individuation is critical to the formation of discrete perceptual representations, individuation follows the initial registration of low-level stimulus features (23). Our account explains why the earliest stimulus-driven responses in V4, occurring 60-100 ms after stimulus onset, show robust increases in amplitude but little or no evidence for earlier latency of responses at attended positions (13–15). Likewise, our account explains why human EEG studies of the P1 component, which occurs 80-130 ms after stimulus onset and is thought to reflect the first feedforward sweep of activity in extrastriate cortex (10, 39), reveal clear increases in amplitude but no change in the latency of responses to stimuli at attended positions (9–12). In contrast to these very early responses, the N2pc component, which is speeded by attention, is a mid-latency component that occurs between 150 and 300 ms after stimulus onset (16, 17). Thus, our findings suggest that attention speeds the formation of discrete object representations without changing the latency of the first wave of sensory encoding.

## Conclusions

We showed that the target-evoked N2pc, a neural marker of object individuation, is speeded for targets that appear at attended locations. This finding provides neural evidence that bolsters the conclusions of past behavioral studies that attention speeds visual processing, while reconciling these findings with work that has not found latency shifts in the earliest visually evoked neural responses. Although attention may not speed the earliest stages of sensory processing, our results suggest that attention does speed the critical transition between raw sensory encoding and the formation of individuated object representations.

## Materials and Methods

### Subjects

Sixty-three volunteers (25 in Experiment 1, and 38 in Experiment 2) participated in the experiments for monetary compensation ($15 per hour). Subjects were between 18 and 35 years old, reported normal or corrected-to-normal visual acuity, and provided informed consent according to procedures approved by the University of Chicago Institutional Review Board.

#### Experiment 1

Our target sample size was 16 subjects in Experiment 1. We excluded subjects that had fewer than 600 artifact-free trials per condition. We excluded a total of nine subjects from our sample. Data collection was terminated early for two subjects because the data was unusable due to excessive artifacts and/or poor task performance. In addition, seven subjects were excluded because of excessive artifacts. Thus, our final sample included 16 subjects (9 male, 7 female; mean age = 22.9 years, *SD* = 2.8). Subjects in the final sample provided 802 trials on average (*SD* = 95) in the informative-cue condition, and 807 trials on average (*SD* = 80) in the uninformative-cue condition.

#### Experiment 2

Our target sample size was 24 subjects in Experiment 2 (50% larger than Experiment 1 to increase statistical power). Again, we excluded subjects that had fewer than 600 artifact-free trials per condition. In addition, we terminated data collection for subjects for whom we could not obtain usable eye tracking data. We excluded a total of 12 subjects from our sample. Data collection was terminated early for eight subjects and these subjects were excluded from our sample for the following reasons: the subject was making too many eye movements during trials (1 subject), we were unable to obtaining usable eye tracking data and the session was running behind schedule (3 subjects), the subject was feeling unwell (1 subject), the subject withdrew from the study (1 subject), the experimenter forgot to save the EEG data (1 subject), and an equipment failure disrupted data collection (1 subject). In addition, four subjects were excluded because of excessive artifacts. Thus, the final sample included 26 subjects (11 male, 15 female; mean age = 22.9 years, *SD* = 3.9). We overshot our target sample size of 24 because the sessions for our final subjects were scheduled before we reached our target sample size. Subjects in the final sample provided 833 trials on average (*SD* = 100) in the informative-cue condition, and 841 trials on average (*SD* = 99) in the uninformative-cue condition.

### Apparatus and Stimuli

We tested the subjects in a dimly lit, electrically shielded chamber. Stimuli were generated using Matlab (The Mathworks, Natick, MA) and the Psychophysics Toolbox (40, 41), and were presented on a 24-in. LCD monitor (refresh rate: 120 Hz, resolution: 1080 × 1920 pixels) at a viewing distance of approximately 79 cm for Experiment 1 and 75 cm for Experiment 2. Stimuli were rendered in dark gray against a medium gray background.

### Task Procedures

#### Experiment 1

Subjects performed a cued-search task [c.f. (22)]. On each trial, they searched for a target (a diamond) among seven distractors (squares), and reported whether the target was missing the left or right corner (Fig. 1). In some blocks, a cue indicated the exact location of the target in advance, which enabled observers to attend the target location prior to the onset of the search array (*informative* cue). In other blocks, the cue provided no information about the location of the impending target (*uninformative* cue).

A fixation point (0.15° in diameter) was present throughout each block of trials. Each trial began with a 100-ms cue. In informative-cue blocks, the cue was a bar (0.125° long, 0.08° wide) that extended from the fixation point, and pointed to the location where the target would appear. In uninformative-cue blocks, the fixation point increased in size (to 0.2°). Thus, uninformative cues provided the same temporal information as informative cues, but provided no information about the target location. After a 600-ms inter-stimulus interval, a search array appeared for 200 ms. Each search array comprised eight stimuli (a target among seven distractors) equally spaced around fixation at an eccentricity of 4° (Fig. 1). The distractors were squares (1.6° × 1.6°). The target was a diamond (identical in size to the distractors) that was missing the left or right corner. Subjects reported which corner by pressing the “z” key (left hand) or the “/” key (right hand). Subjects were instructed to respond as quickly as possible while maintaining high accuracy. Each trial was terminated by the subject’s response, and was followed by a variable inter-trial interval (ITI) between 1500 and 1800 ms. Subjects were provided with feedback about their performance (mean response time and accuracy) at the end of each block of trials. To minimize ocular artifacts during the trials, we instructed subjects to maintain fixation throughout each block of trials, and to blink shortly after their response, before the next trial began.

Subjects completed 32 blocks of 64 trials, with two exceptions: one who completed 30 blocks, and another who completed 27 blocks. Within each block, the target appeared at each of the eight possible locations equally often. The cue conditions (informative or uninformative) were alternated across blocks, and the order of conditions (informative or uninformative first) was counterbalanced across subjects. Subjects completed between two and four practice blocks (as needed based on task performance).

#### Experiment 2

Subjects completed the cued-search task used in Experiment 1 with three changes. First, we used a different cue: eight bars extended from the fixation point (0.125° long, 0.08° wide), one pointing towards each of the eight stimulus positions. In informative-cue blocks, one bar that was different color to the rest (either a red bar among blue bars or a blue bar among red bars, counterbalanced across subjects) pointed toward the target location. In uninformative-cue blocks, all bars were the same color (blue or red, counterbalanced across subjects). The red and blue colors used for the cues were closely matched for luminance. Second, we used a longer ITI (jittered between 1500 ms and 1900 ms). Third, to cue subjects to blink during the ITI, the fixation point disappeared 200 ms after the subject made their response (i.e., 200 ms after the ITI began), and re-appeared 500-600 ms before the cue for the next trial. We instructed subjects to blink when the fixation point was absent. Finally, we recalibrated they eye tracker in between trials if necessary. After calibration, the fixation point was presented for 500 ms before starting the next trial.

Subjects completed 32 blocks except for two subjects who completed 30 blocks, one subject who completed 24 blocks, and another subject who completed 36 blocks. For the subject who completed 36 blocks, EEG data was not recorded for 5 blocks due to experimenter error (see Glitches), so data were analyzed only for the 31 blocks with EEG data.

### EEG Acquisition

We recorded EEG activity using 30 active Ag/AgCl electrodes mounted in an elastic cap (Brain Products actiCHamp, Munich, Germany). We recorded from International 10/20 sites Fp1, Fp2, F7, F3, Fz, F4, F8, FC5, FC1, FC2, FC6, C3, Cz, C4, CP5, CP1, CP2, CP6, P7, P3, Pz, P4, P8, PO7, PO3, PO4, PO8, O1, Oz, and O2. Two additional electrodes were placed on the left and right mastoids, and a ground electrode was placed at position Fpz. All sites were recorded with a right-mastoid reference, and were re-referenced offline to the algebraic average of the left and right mastoids. We recorded electrooculogram (EOG) with passive electrodes, with a ground electrode placed on the left check. Horizontal EOG was recorded with a bipolar pair of electrodes placed ~1cm from the external canthus of each eye, and vertical EOG with a bipolar pair of electrodes placed above and below the right eye. Data were filtered online (low cut-off = 0.01 Hz, high cut-off = 80 Hz, slope from low-to-high cut-off = 12 dB/octave), and were digitized at 500 Hz using BrainVision Recorder (Brain Products, Munich, Germany) running on a PC. We maintained impedances below 10 kΩ.

### Eye Tracking

We recorded gaze position using a desk-mounted infrared eye-tracking system (EyeLink 1000 Plus, SR Research, Ontario, Canada). According to the manufacturer, this system provides spatial resolution of 0.01° of visual angle, and average accuracy of 0.25-0.50° of visual angle. Gaze position was sampled at 1000 Hz. Stable head position was maintained during the task using a chin rest. The eye tracker was re-calibrated as needed throughout the session, including whenever subject removed their chin from the chin rest.

### Artifact Rejection

We used a semi-automated procedure to remove trials that where contaminated by EEG artifacts (amplifier saturation, excessive muscle noise, skin potentials) or by ocular artifacts (blinks and eye movements). After an initial automated routine to identify trials that contained artifacts, the segmented data were visually inspected. We excluded trials contaminated by artifacts from all analyses (including behavioral analyses). We discarded electrodes Fp1 and Fp2 for all subjects because data quality was often poor (i.e. excessive high-frequency noise or slow drifts) at these sites. Furthermore, we discarded some data from one or two additional electrodes for some subjects because of low quality data (excessive high-frequency noise, drifts, or sudden steps in voltage). Subjects were excluded from the final sample if they had fewer than 600 artifact-free trials per condition (see Subjects).

### N2pc analysis

For trials with correct responses, we measured the target-evoked N2pc locked to the search array by calculating a contralateral-ipsilateral difference wave averaging across posterior electrodes P7/8, PO7/8, PO3/4 and O1/2. For one subject, EEG data was unusable for electrode P7, therefore we measured the N2pc at the electrode PO7/8, PO3/4, and O1/2. We baseline corrected the waveforms by subtracting the mean voltage in the 200-ms period prior to onset of the search array.

We used a jackknife procedure (42) to test for differences in the onset latency of the N2pc following informative and uninformative cues. N2pc onset latency was measured at the earliest time at which the N2pc difference wave reached 50% of its maximum amplitude. The latency difference between conditions, *D*, was measured as the difference in onset latency between conditions in the subject-averaged N2pc difference waves. We used a jackknife procedure to estimate the standard error of the latency difference, *SE*_*D*_, from the latency differences obtained for subsamples that included all but one subject (42). Specifically, the latency differences, *D*_−*i*_ (for *i* = 1, …, *N*, where *N* is the sample size), were calculated where *D*_−*i*_ was the latency difference for the sample with all subjects except for subject *i*. The jackknife estimate of the *SE*_*D*_ was calculated as

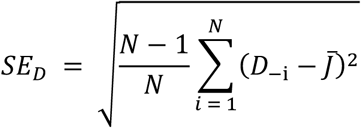

where 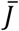 is the mean of the differences obtained for all subsamples (i.e., 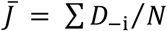. A jackknifed *t* statistic, *t*_*j*_, was then calculated as

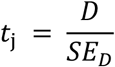

which follows an approximate *t* distribution with *N* – 1 degrees of freedom under the null hypothesis. These tests were one-tailed because we had the clear directional hypothesis that N2pc onset would be earlier following informative cues than following uninformative cues.

### Time-frequency Analysis

To calculate alpha-band power at each electrode, we first band-pass filtered the raw EEG data using EEGLAB between 8 and 12 Hz [“eegfilt.m” (43)]. We applied a Hilbert transform (Matlab Signal Processing Toolbox) to the band-pass–filtered data to obtain the complex analytic signal. Instantaneous power was calculated by squaring the complex magnitude of the complex analytic signal. We used a 375-ms long filter kernel (i.e., 3 cycles with a low cutoff of 8 Hz). Thus, blurring in the time domain due to filtering did not extend beyond 188 ms before or after each time point.

### Inverted Encoding Model

Following our past work (31, 37), we used an inverted encoding model (44–46) to reconstruct spatially selective channel-tuning functions (CTFs) from the pattern of alpha-band (8–12 Hz) power across electrodes. This analysis assumed that alpha power at each electrode reflects the weighted sum of eight spatially selective channels (i.e., neuronal populations), each tuned for a different position in the visual field. Specifically, we modeled alpha power at each electrode as the weighted sum of eight spatial channels tuned for eight locations equally spaced around the central fixation point corresponding to positions the stimuli in the search array appeared at (Fig. 1). We modeled the response profile of each spatial channel across angular locations as a half sinusoid raised to the twenty-fifth power:

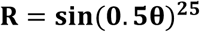

where θ is angular location (0–359°), and *R* is the response of the spatial channel in arbitrary units. We circularly shifted this basis function to obtains a basis set of eight basis function tuned for the eight equally spaced angular locations (0°, 45°, 90°, etc.).

An IEM analysis was applied to each time point to obtain time-resolved CTFs. We partitioned our data into independent sets of training data and test data (see the Training and Test Data section). The analysis proceeded in two stages (training and test). In the training stage, training data (*B*_*1*_) were used to estimate weights that approximate the relative contribution of the eight spatial channels to the observed response measured at each electrode. Let *B*_*1*_ (*m* electrodes × *n*_*1*_ measurements) be the power at each electrode for each measurement in the training set, *C*_*1*_ (*k* channels × *n*_*1*_ measurements) be the predicted response of each spatial channel (determined by the basis functions) for each measurement, and *W* (*m* electrodes × *k* channels) be a weight matrix that characterizes a linear mapping from “channel space” to “electrode space”. The relationship between *B*_*1*_, *C*_*1*_, and *W* can be described by a general linear model of the form:

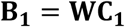

The weight matrix was obtained via least-squares estimation as follows:

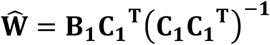

In the test stage, we inverted the model to transform the independent test data *B*_*2*_ (*m* electrodes × *n*_*2*_ measurements) into estimated channel responses, *C*_*2*_ (*k* channels × *n*_*2*_ measurements), using the estimated weight matrix, *Ŵ*, that we obtained in the training phase:

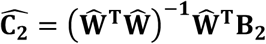

Each estimated channel response function was then circularly shifted to a common center, so the center channel was the channel tuned for the position of the cued/target location (a Channel Offset of 0°), then averaged these shifted channel-response functions across the eight position bins to obtain a CTF. The IEM analysis was applied to each subject separately because the exact contributions of each spatial channel to each electrode (i.e., the channel weights, *W*) will likely vary across individuals.

### Training and test data

For the IEM procedure, we partitioned artifact-free trials into independent sets of training data and test data for each subject. When comparing CTF properties across conditions, it is important to estimate a single encoding model that is then used to reconstruct CTFs for each condition separately. If this condition is not met, then it is difficult to interpret differences in CTF selectivity between conditions because these might result from differences between the training sets [for further discussion of this issue, see (47)]. Here, we used data from the informative-cue condition to estimate the encoding model in the training phase, and we reconstructed CTFs for each of the condition (informative cue and uninformative cue) separately. Specifically, we partitioned data in each condition into three independent sets, equating the number of trials for each location within each of the six sets (three informative cue sets and three uninformative cue sets). For each set, we averaged across trials for each stimulus location bin to obtain estimates of alpha power values across all electrodes for each target location (electrodes × locations, for each time point). We used a leave-one-out cross-validation routine such that two of the three sets of informative-cue data served as the training data. The remaining set of informative-cue data served as the test data for that condition, and one of the sets of the data from the uninformative-cue condition serves as the test data for that condition. We applied the IEM routine using each of the three matrices within each condition as the data, and the remaining two informative-cue sets as the training set. The resulting CTFs were averaged across the three test sets for each condition.

Because we equated the number of trials for each target location within each set of trials, some trials were not assigned to any set. Thus, we used an iterative approach to make use of all available trials. For each iteration, we randomly partitioned the trials into six sets (as just described), and performed the IEM procedure on the resulting training and test data (such that trials that were not included in any block varied across iterations), and we averaged the resulting channel-response profiles across iterations. We performed a total of 50 iterations per subject.

### CTF selectivity

To quantify the spatial selectivity of alpha-band CTFs, we used linear regression to estimate CTF slope (31, 37). Specifically, we calculated the slope of the channel responses as a function of spatial channels after collapsing across channels that were equidistant from the channel tuned for the position of the stimulus (i.e., a channel offset of 0°). Higher CTF slope indicates greater spatial selectivity.

### Cluster-based permutation test

We used a cluster-based permutation test to identify when CTF selectivity was reliably above chance, which corrects for multiple comparisons (48, 49). We identified clusters in which CTF selectivity was greater than zero by performing a one-sample *t*-test (against zero) at each time point. We then identified clusters of contiguous points that exceeded a *t*-statistic threshold corresponding to a one-side *p*-value of .05. For each cluster, we calculated a test statistic by summing all *t-values* in the cluster. We used a Monte Carlo randomization procedure to empirically approximate a null-distribution for this test statistic. Specifically, we repeated the IEM procedure 1000 times but randomized the position labels within each training/test set, such that the labels were random with respect to the observed response at each electrode. For each run of the analysis with randomized position labels, we identified clusters as described above, and recorded the largest test statistic, resulting in a null distribution of 1000 cluster test-statistics. With this null distribution in hand, we identified clusters in our unpermuted data that had test statistics larger than the 95_th_ percentile of the null distribution. Thus, our cluster test was a one-tailed test with an alpha level of .05, corrected for multiple comparisons.

### Glitches

#### Experiment 1

For seven subjects, eye tracking data was not recorded for a subset of trials due to a glitch with the eye tracker. Five of these subjects were included in our final sample. For these subjects, between 60 and 240 of the trials that remained after exclusion of trials with artifacts and incorrect responses were missing eye tracking data.

For one subject, the stimulus presentation computer crashed during one block of the task. We excluded data from this block from all analyses.

For another subject, we failed to record EEG data for 25 trials due to an equipment failure. We excluded these trials from all analyses.

#### Experiment 2

For a subset of subjects, the EOG electrodes were plugged in incorrectly. This problem does not affect our analyses because we relied exclusively on eye tracking data for detection of ocular artifacts and analyses of gaze position.

For one subject, the experimenter forgot to resuming saving EEG data after a break between blocks. Thus, we are missing EEG data for 5 blocks (320 trials). We excluded these blocks from all analysis. This subject completed a total of 36 blocks. Thus, we have 31 usable blocks of data for this subject.

We terminated data collection for one subject because of technical difficulties with the EEG amplifier.

## Data and Code Availability

All data and code will be made available on Open Science Framework at the time of publication.

## Acknowledgements

This work was supported by National Institute of Mental Health Grant 5RO1 MH087214-08. We thank Mei Arditi, Ariana Gale, and Russell Jaffe for assistance with data collection.

## SUPPLEMENTARY INFORMATION

### The effect of spatial cuing on N2pc latency cannot be explained by residual biases in gaze position

We recorded eye position using an infrared eye tracker, and used a stringent threshold for rejecting trials with blinks or eye movements (see Materials and Methods in the main text). Nevertheless, we found very small biases in eye position toward the cued location following informative cues remained after artifact rejection. Figure S1 shows mean gaze position during the search array (700-900 ms after cue onset) as a function of target position following informative and uninformative cues.^1^ Following informative cues, mean gaze position varied by less than 0.15° (approximately the size of our fixation point). Thus, while there was a detectable bias in gaze position toward the cued location, this bias was very small as would be expected after artifact rejection. Following uninformative cues, no such variation in gaze position was seen.

As a result of this small bias in gaze position, the target stimulus in the search display appeared marginally closer to fovea on average following informative cues that following uninformative cues. Thus, we tested whether this very small bias in eye position toward the target location could explain the earlier onset of the N2pc component following informative cues than following uninformative cues (Fig. 3). We performed this analysis using data from Experiment 2, in which we obtained reliable eye tracking data for all subjects. For each subject, we median split trials on the basis of the distance between mean gaze position (during the search array) and the target location. Thus, trials were categorized as having gaze position *biased-toward* (the 50% of trials in which gaze position was closest to the target location) or *biased-away* (the 50% of trials in which the gaze position was farthest from the target location) from the target location. Note that this sorting of trials based on eye position is relative: a trial that was categorized as “biased-toward” does not necessarily imply that the gaze position was to the target-side of the fixation dot for that trial. We sorted trials based on gaze position for each cue condition (informative and uninformative) separately. Figure S2 shows the mean gaze coordinates for the bias-toward and bias-away trials for each cue condition. This plot shows that sorting the trials based on gaze position created substantial bias in gaze position toward or away from the target position.

To test whether bias in gaze position influences the latency of the N2pc component, we examined the N2pc as a function of gaze position. Figure S3 shows the N2pc difference waves for biased-toward and biased-away trials for each condition. We used a jackknife-based procedure to test for differences in the latency of the N2pc component (see Materials and Methods in the main text). We found that the N2pc did not onset earlier for biased toward than for biased away trials for either the informative-cue condition, *t*(25) = −0.74, *p* = 0.77 (one-tailed), or for the uninformative-cue condition, *t*(25) = 0.0, *p* = 0.5 (one-tailed). However, a robust latency different persisted between cue conditions for both biased-toward trials, *t*(25) = 3.50, *p* < .001 (one-sided), and biased-away trials, *t*(25) = 4.05, *p* < .001 (one-tailed). Thus, the effect of spatial cues on the latency of the N2pc component cannot be explained by small residual biases in eye position that persist after artifact rejection.

**Figure S1.**
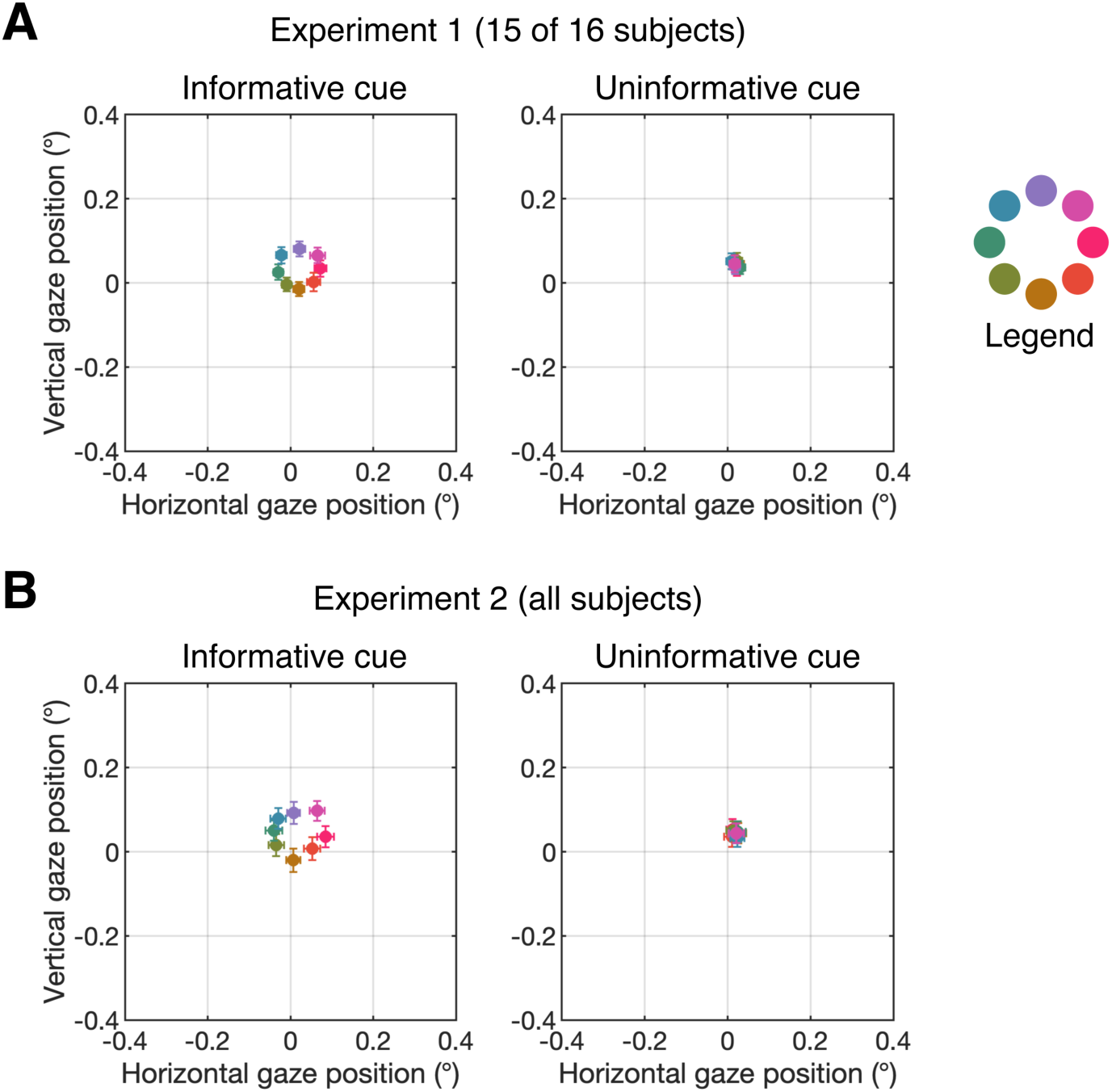
Mean gaze position during the search array following informative cues (left) and uninformative cues (right) for Experiment 1 (A) and Experiment 2 (B). The legend at the right of the plot shows which color corresponds to each of the eight target positions. Error bars show ±1 SEM across subjects.

**Figure S2.**
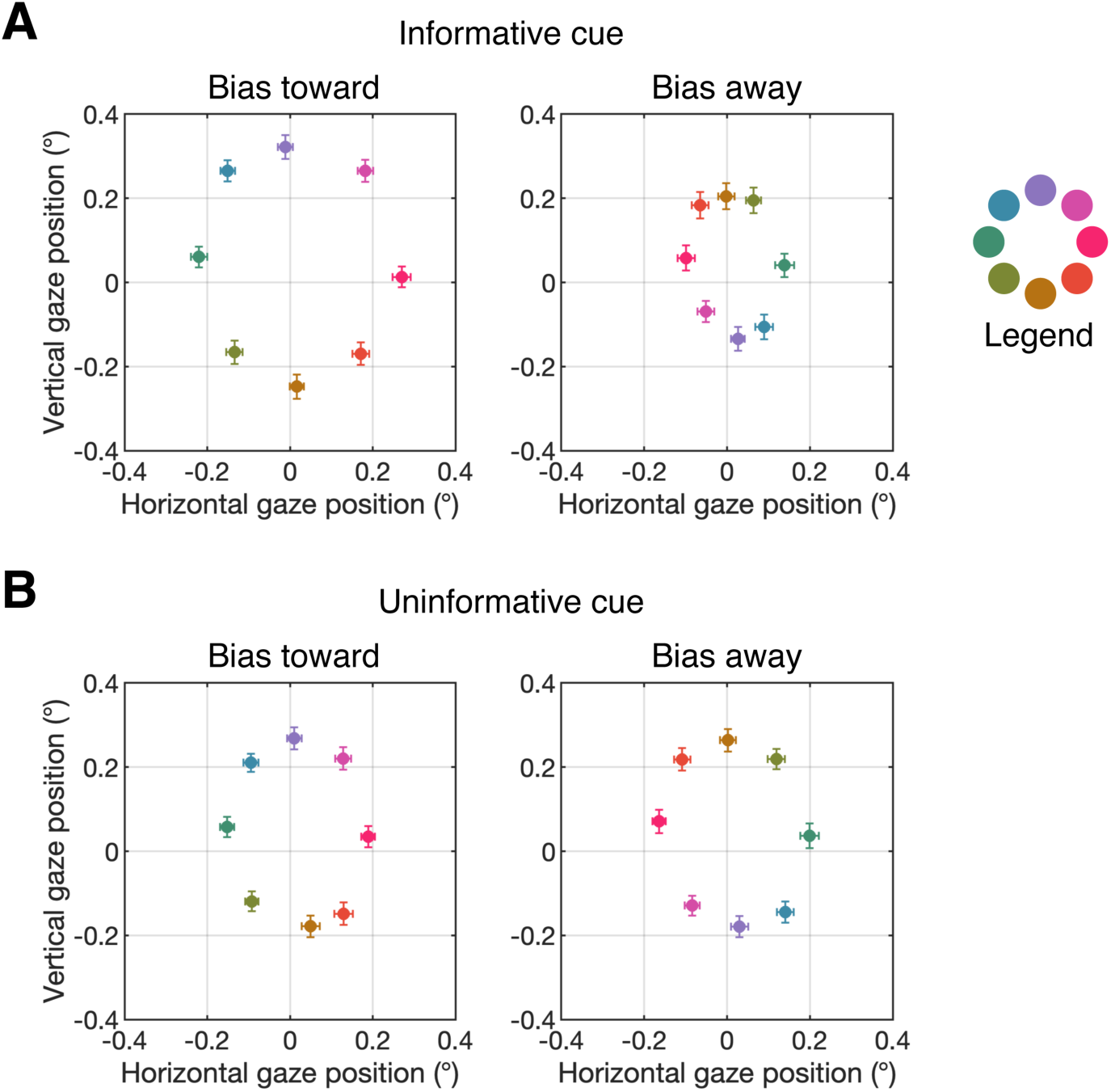
Mean gaze position during the search array for bias-toward (left) and bias-away (right) trials (right) following informative cues (A) and uninformative cues (B). The legend at the right of the plot shows which color corresponds to each of the eight target positions. Error bars show ±1 SEM across subjects.

**Figure S3.**
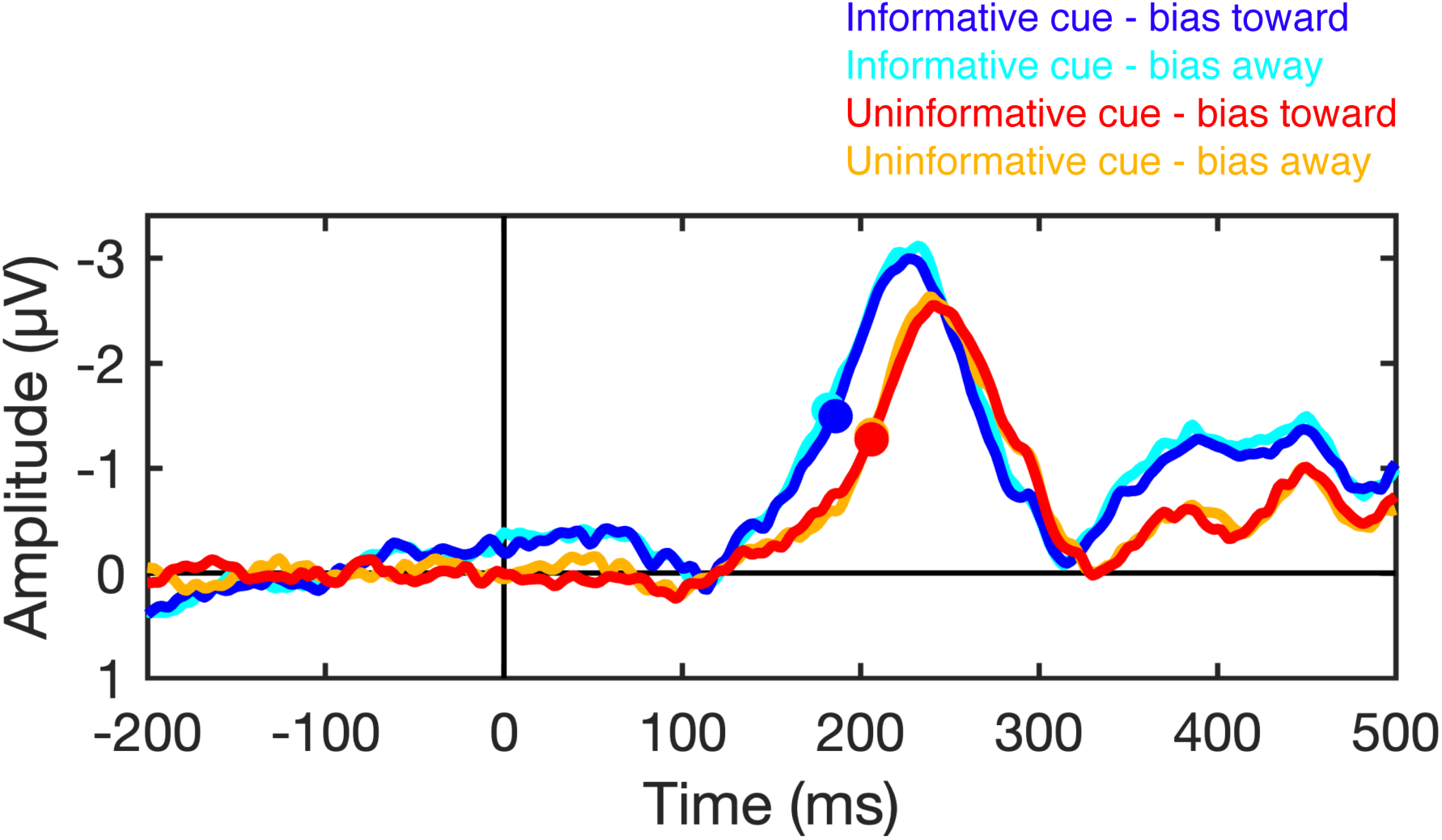
N2pc latency is not sensitive to residual variation in gaze position. The plots show the N2pc difference waves (contralateral minus ipsilateral) time locked to the onset of the search array when gaze position was biased toward or biased away from the target location following informative and uninformative cues. The filled circles mark the time at which the N2pc reached the onset criterion (50% of maximum amplitude) for each condition.

## Footnotes

1 For Experiment 1, we were unable to obtain usable eye tracking data for one subject. Furthermore, between 60 and 240 trials (after artifact rejection) were missing eye tracking data for five subjects due to a bug with the eye tracker (see Materials and Methods, Glitches), which were omitted from the analysis of gaze position.

